# Increased fatty acid metabolism and decreased glycolysis are hallmarks of metabolic reprogramming in the brain during recovery from experimental stroke

**DOI:** 10.1101/2022.03.22.485395

**Authors:** Sanna H. Loppi, Marco A. Tavera-Garcia, Danielle A. Becktel, Boaz K. Maiyo, Kristos E. Johnson, Rick G. Schnellmann, Kristian P. Doyle

## Abstract

The goal of this study was to evaluate changes in metabolic homeostasis during the first 12 weeks of recovery in a distal middle cerebral artery occlusion mouse model of stroke. To achieve this goal, we compared the brain metabolomes of ipsilateral and contralateral hemispheres from aged male mice up to 12 weeks after stroke to that of age-matched naïve and sham operated mice. There were 707 biochemicals detected in each sample by liquid chromatography-mass spectroscopy (LC-MS). Mitochondrial fatty acid β-oxidation, indicated by acyl carnitine levels, was increased in stroked tissue at 1 day and 4 weeks following stroke. Glucose and several glycolytic intermediates were elevated in the ipsilateral hemisphere for 12 weeks compared to the aged naïve controls, but pyruvate was decreased. Additionally, itaconate, a glycolysis inhibitor associated with activation of anti-inflammatory mechanisms in myeloid cells, was higher in the same comparisons. These changes correlated with reduced levels of glutamate, dopamine, and adenosine in the ipsilateral hemisphere after stroke. These results indicate that chronic metabolic differences exist between stroked and control tissue, including alterations in fatty acid metabolism and glycolysis for at least 12 weeks after stroke.

## Introduction

Stroke is among the leading causes of death and disability, and markedly diminishes the quality of life of stroke survivors.^1^ While mechanical thrombectomy can be used to treat large vessel occlusions in the anterior circulation,^2^ recombinant tissue plasminogen activator (r-tPA) remains the only clinically approved pharmacological treatment for ischemic stroke. However, r-tPA is only suitable for a fraction of patients due to its side-effects and narrow therapeutic time window.^3^ There is an urgent need to identify new targets for improving stroke recovery.

Ischemic stroke is caused by the blockage of a brain blood vessel which results in insufficient delivery of oxygen and glucose to support cellular homeostasis. Although this acute perturbation in brain metabolism is well characterized, little is known about how brain metabolism is altered in the weeks and months during recovery. This is an important knowledge gap to address because although cells adapt to their environment by undergoing metabolic reprogramming, metabolic reprogramming alters cellular function.^4^

Therefore, the goal of this study was to evaluate how metabolism is altered in the brain in the first 12 weeks after stroke, which is when most recovery takes place.^5^ To accomplish this goal, we performed liquid chromatography-mass spectroscopy (LC-MS) global metabolomics on brain tissue from 7 month and 18– to 20-month-old C57BL/6N mice to determine how aging impacts brain metabolism as most strokes occur in people over the age of 65.^6^

We then performed LC-MS global metabolomics on 18– to 20-month-old mice sacrificed at 1 day and 2, 4, 8 and 12 weeks after stroke or sham surgery to evaluate how stroke impacts brain metabolism during recovery. Data from the contralateral and ipsilateral hemispheres were compared to determine if changes in metabolism were global or region specific.

## Material and Methods

### Animals

Sample sizes for the study were determined using power analyses based on expected variances and group differences. A total of 82 male C57BL/6N mice were acquired from the National Institute on Aging. Six of them were young adults (7-month-old), and 76 were aged (18- to 20- month-old). The young mice were sacrificed without undergoing stroke or sham surgery (young naïve group). The aged mice were randomly divided into the following three groups: aged naïve (*n* = 6), aged sham (*n* = 30), and aged stroke (*n* = 40). The aged stroke and sham mice were sacrificed at 1 day post ischemia (dpi) (7 stroke and 6 sham) and 2 weeks (7 stroke and 6 sham), 4 weeks (7 stroke and 6 sham), 8 weeks (6 stroke and 6 sham), and 12 weeks (6 stroke and 6 sham) after stroke or sham surgery (Figure 1A). The pre-set exclusion criteria of the study were (1) unsuccessful induction of ischemia (4 mice), (2) death of the animal during the experiment (3 mice), and (3) being a statistically significant outlier in any of the analyses (4 mice). All animal experiments followed the NIH guidelines and were approved by the University of Arizona Institutional Animal Care and Use Committee. RIGOR criteria and ARRIVE guidelines were followed when conducting and reporting the experiments.^7,8^ Every effort was made to minimize the harm and suffering of the animals.

**Figure 1.**
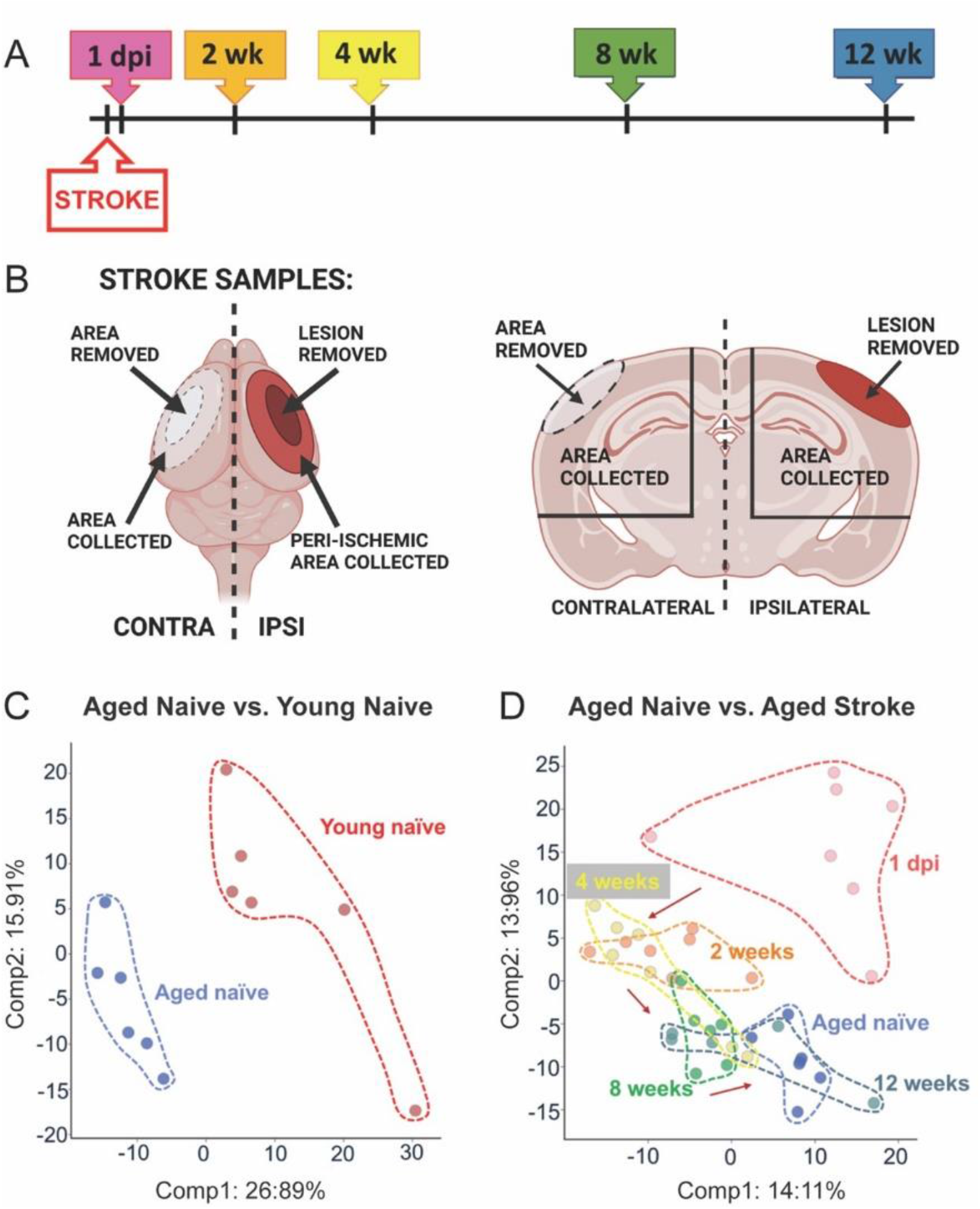
Study design and principal component analysis. **A.** Timeline indicating the time points of tissue collection after stroke. **B.** Schematic illustration of the locations where the samples were collected (created with BioRender.com). **C.** Principal component analysis showing a clear separation between the young and the aged naïve mice along Principal Component 1. **D.** Principal component analysis between the aged naïve mice and the stroke groups showing separation based on stroke time point.

### Stroke and sham surgeries

The permanent distal middle cerebral artery (MCA) occlusion was carried out as described previously.^9^ Briefly, anesthesia was induced with 3% isoflurane (JD Medical, Phoenix, AZ, USA), and maintained throughout surgery. After the temporalis muscle was exposed and retracted, a hole (1 mm in diameter) was drilled into the temporal bone to expose the MCA. The dura was removed, after which the artery was cauterized. The temporalis muscle was replaced, and the wound closed using surgical glue. Body temperature of the animals was maintained at 37 ± 0.5 °C during the procedure using a heating pad equipped with a rectal probe. Respiration and temperature were monitored throughout surgery. Immediately after surgery, mice were placed in a hypoxia chamber (Coy Laboratory products, Grass Lake, MI, USA) containing 9% oxygen and 91 % nitrogen for 45 minutes. The purpose of hypoxia in this model is to increase infarct size and reduce variability in infarct size.^9^ A single dose of buprenorphine hydrochloride (Buprenex® Injection 0.3 mg/mL, Henry Schein Medical, Melville, NY, USA; 0.1 mg/kg s.c.) was administered prior to surgery, and sustained release buprenorphine (Buprenorphine SR 1 mg/mL, Zoopharm LLC, Laramie, WY, USA; 1 mg/kg s.c.) was administered 24 hours after surgery as a post-operative analgesic. Sham operated mice went through the same procedure, except for cauterization of the MCA.

### Perfusion and tissue collection

At the time points described in Figure 1A, mice were anesthetized with 3% isoflurane. Blood samples were collected from the heart, after which mice were transcardially perfused with 0.9% saline solution. Infarcts and the corresponding regions from sham operated mice were dissected and removed. The corresponding areas of the contralateral hemisphere were also dissected and removed. After removal of the olfactory bulb and cerebellum, the remaining ipsilateral and contralateral hemispheres were snap frozen in liquid nitrogen and shipped to Metabolon Inc. (Morrisville, NC, USA) for metabolomic analysis. The tissue was not further separated into smaller brain regions due to a minimum requirement of 50mg of tissue outlined in Metabolon’s global metabolic profiling protocol. A guide to the dissection is provided in Figure 1B.

### Metabolomic analysis

Global metabolite profiling analysis on brain tissue was performed by Metabolon Inc. using ultra high-performance liquid chromatography coupled to tandem mass spectrometry (UPLC-MS/MS). In brief, following receipt by Metabolon, samples were inventoried and immediately stored at −80°C. Samples were prepared using the automated MicroLab STAR® system from Hamilton Company (Reno, NV/Franklin, MA, USA). Several recovery standards were added prior to the first step in the extraction process for quality control (QC) purposes. To remove protein, dissociate small molecules bound to protein or trapped in the precipitated protein matrix, and to recover chemically diverse metabolites, proteins were precipitated with methanol under vigorous shaking for 2 min (GenoGrinder 2000; Glen Mills, Clifton, NJ, USA) followed by centrifugation. The resulting extract was divided into five fractions: two for analysis by two separate reverse phase (RP)/UPLC-MS/MS methods with positive ion mode electrospray ionization (ESI), one for analysis by RP/UPLC-MS/MS with negative ion mode ESI, one for analysis by HILIC/UPLC-MS/MS with negative ion mode ESI, and one reserved for backup. Samples were placed briefly on a TurboVap® (Zymark, Biotage, Uppsala, Sweden) to remove the organic solvent. The sample extracts were stored overnight under nitrogen before preparation for analysis. Several types of controls were analyzed in concert with the experimental samples: a pooled matrix sample generated by taking a small volume of each experimental sample served as a technical replicate throughout the data set; extracted water samples served as process blanks; and a cocktail of QC standards that were carefully chosen not to interfere with the measurement of endogenous compounds were spiked into every analyzed sample, allowed instrument performance monitoring, and aided chromatographic alignment.

Instrument variability was determined by calculating the median relative standard deviation (RSD) for the standards that were added to each sample prior to injection into the mass spectrometers. Overall process variability was determined by calculating the median RSD for all endogenous metabolites (i.e., non-instrument standards) present in 100% of the pooled matrix samples. Experimental samples were randomized across the platform run with QC samples spaced evenly.

For UPLC-MS/MS, all methods utilized a Waters ACQUITY ultra-performance liquid chromatography (UPLC) and a Thermo Scientific Q-Exactive high resolution/accurate mass spectrometer interfaced with a heated electrospray ionization (HESI-II) source and Orbitrap mass analyzer operated at 35,000 mass resolution. The sample extract was dried then reconstituted in solvents compatible to each of the four methods.

For data extraction and compound identification, raw data was extracted, peak-identified and QC processed using Metabolon’s hardware and software. Compounds were identified by comparison to library entries of purified standards or recurrent unknown entities. Metabolon maintains a library based on authenticated standards that contains the retention time/index (RI), mass to charge ratio (m/z), and chromatographic data (including MS/MS spectral data) on all molecules present in the library. Furthermore, biochemical identifications were based on three criteria: retention index within a narrow RI window of the proposed identification, accurate mass match to the library +/− 10 ppm, and the MS/MS forward and reverse scores between the experimental data and authentic standards.

### Neurofilament light (NF-L) assay

To assess the level of neurodegeneration, plasma samples were sent to PBL Assay Science (Piscataway, NJ, USA) and analyzed using the Simoa™ NF-Light® kit (Cat #103186, Quanterix, Billerica, MA, USA).

### Statistics

To generate the heat maps, Welch’s two-sample *t*-test was used to test whether two unknown means were different from two independent populations. To generate the bar graphs significance was tested using an unpaired two-tailed *t* test with Welch’s correction, or Brown-Forsythe and Welch’s ANOVA test, followed by Dunnett’s T3 multiple comparisons test, as indicated in the figure legends. For principal component analysis, each principal component is a linear combination of every metabolite, and the principal components are uncorrelated. The first principal component was computed by determining the coefficients of the metabolites that maximized the variance of the linear combination. The second component finds the coefficients that maximize the variance with the condition that the second component is orthogonal to the first. Statistical analyses were performed using GraphPad Prism software 9.3.1 (GraphPad Software, LaJolla, CA, USA), and normality was assessed using the Kolmogorov-Smirnov test. Statistical tests and sample sizes are provided in each figure legend, and *p* values less than 0.05 were considered to be statistically significant. Statistically significant outliers, calculated using GraphPad Prism QuickCalcs, were excluded from the datasets. Data are presented as mean ± standard deviation (SD).

## Results

### Overview of the analysis

The timeline of the experiment is presented in **Figure 1A**. At each time point, the infarct and corresponding region of the cortex from the contralateral hemisphere were dissected and discarded. The remaining ipsilateral hemisphere, and the equivalent region from the remaining contralateral hemisphere, were then dissected and used for metabolomic analysis (**Figure 1B**). Equivalent brain regions from young naïve mice, aged naïve mice, and sham operated mice were dissected and used for metabolomic analysis as reference controls.

The total number of biochemicals detected by metabolomic analysis of each brain region was 707. At a significance level of *p* < 0.05 (5% of all detected metabolites), 35 differences between groups can be expected by random chance.

**Supplementary Table 1** shows the number of significantly different metabolites when the following comparisons were made: 1) young naïve mice compared to aged naïve mice, 2) stroked mice at each time point compared to aged naïve mice, 3) ipsilateral hemispheres compared to contralateral hemispheres at matched time points, 4) stroked mice compared to sham mice at matched time points, 5) contralateral hemispheres at each time point compared to aged naïve mice, 6) sham mice at each time point compared to aged naïve mice, and 7) contralateral hemispheres compared to sham mice at matched time points. **Supplementary Table 2** provides a full list of the metabolites changed with each comparison.

There were 190 statistically significant changes between the samples from the young and aged naïve mice, while there were 202 changes at 1 dpi when compared to aged naïve brains, 153 changes at 1 dpi when compared to time point-matched contralateral hemispheres, and 167 changes at 1 dpi when compared to time point-matched samples from sham mice.

Although tissue from sham mice and contralateral hemispheres from stroked mice are valid controls for preclinical stroke experiments, there were differences in the metabolic changes in the tissue taken from the aged naïve mice, sham mice, and contralateral hemispheres of the stroked mice. The data in **Supplementary Table 2** provides a reference list of the changes for the sham mice and contralateral hemispheres of the stroked mice when compared to age-matched naïve mice and stroked mice. However, despite these differences, the overall pattern of metabolic changes caused by stroke was comparable regardless of which control was used (**Supplementary Figure 1**).

The difference between young and aged naïve mice was visualized using principal component analysis (PCA) (**Figure 1C**). The PCA plot shows a clear separation of the young and aged mice along Principal Component 1. PCA comparing the aged naïve mice to the stroked mice showed distinct separation based on time point (**Figure 1D**). As expected, the samples collected at 1 dpi showed the most separation, with the later time points clustering progressively closer to the aged naïve controls.

### Metabolic changes caused by aging

Of the 190 identified biochemicals that differed between the brains of young and aged naïve mice, the most prominent alteration was a decrease in the levels of long chain fatty acids in the aged naïve mice (**Figure 2A**). Specifically, there was a reduction in stearate (18:0), oleate/vaccenate (18:1), eicosapentaenoate (EPA; 20:5n3), and docosahexaenoate (DHA; 22:6n3) in the brains of aged naïve mice compared with young naïve mice (**Figures 2B-E)**. Multiple acyl carnitines were increased in aged naïve brains (**Figure 2F**), with myristoylcarnitine (C14) and palmitoleoylcarnitine (C16:1) shown in **Figures 2G-H**. The modification of fatty acids to acyl carnitines is essential for transfer across the inner mitochondrial membrane and subsequent β-oxidation (**Figure 2I**). These data demonstrate that there is an increase in fatty acid metabolism in the aging mouse brain. These results complement previous data indicating that aging-induced mitochondrial dysfunction activates mechanisms for the catabolism of myelin lipids to generate ketone bodies for ATP production.^10^

**Figure 2.**
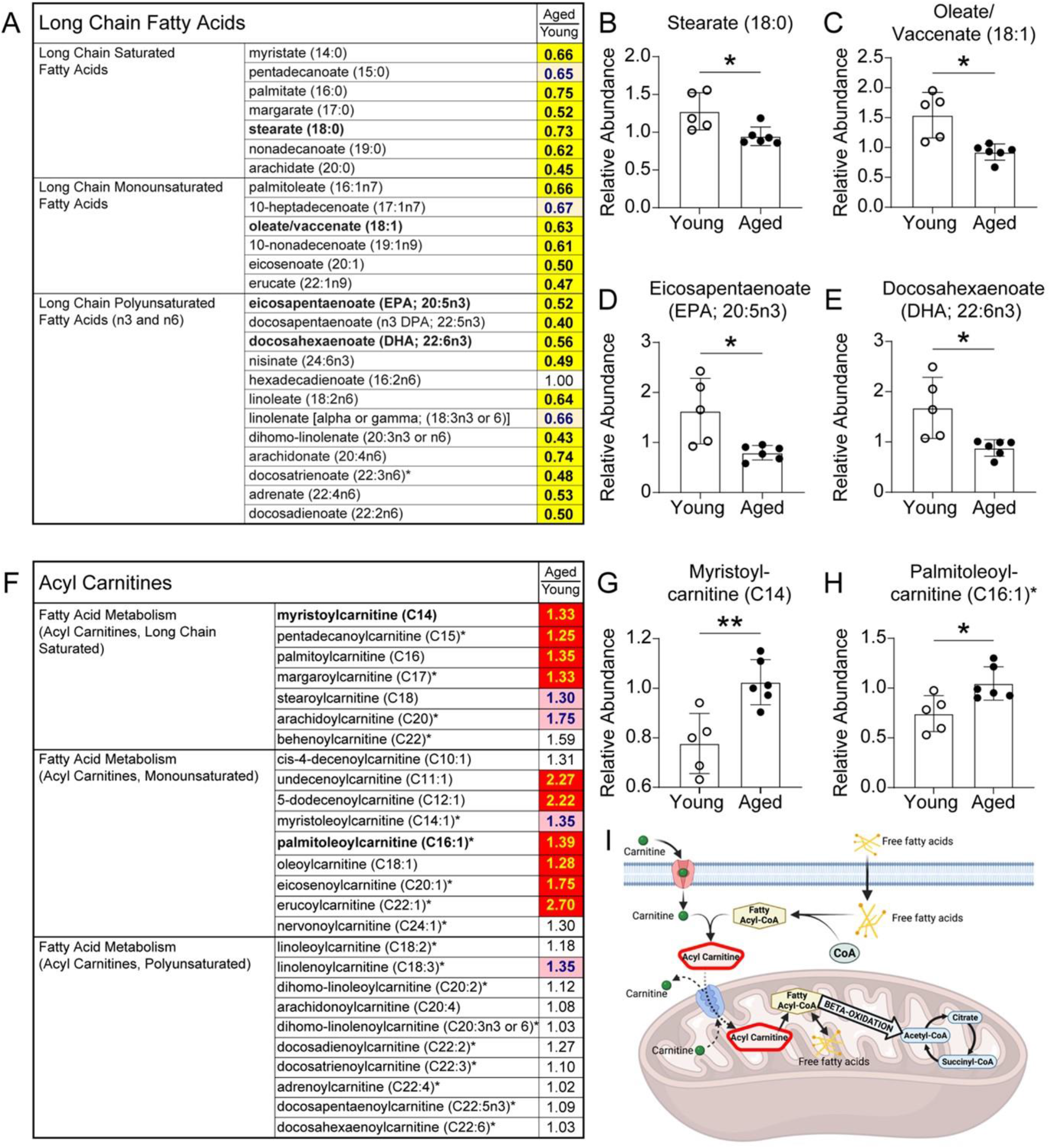
There is a reduction in long chain fatty acids and an increase in acyl carnitines in the aging brain. **A.** Long chain fatty acids were predominantly decreased in aged naïve mice compared to young naïve mice. **B-E.** Representative long chain fatty acids with groups compared by an unpaired, two-tailed t test with Welch’s correction. **B.** Stearate (18:0); *p = 0.0366. **C.** Oleate/vaccenate (18:1); *p = 0.0194. **D.** Eicosapentaenoate (EPA; 20:5n3); *p = 0.0456. **E.** Docosahexaenoate (DHA; 22:6n3); *p = 0.0406. **F.** Some, but not all, species of the medium and long chain acyl carnitines were increased in aged naïve mice compared to young naïve mice. **G-H.** Representative acyl carnitines with groups compared by an unpaired, two-tailed t test with Welch’s correction. **G.** Myristoylcarnitine (C14); **p = 0.0065. H. Palmitoleoylcarnitine (C16:1); *p = 0.0204. n = 5-6. A&F. The numbers in the heatmaps indicate fold change with significance tested by Welch’s two-sample t test. Yellow- and red-shaded cells indicate p ≤ 0.05 (red indicates that the mass-normalized mean values are significantly higher in the condition above the line, and yellow indicates that they are significantly lower). Light red- and light yellow shaded cells indicate 0.05 < p < 0.10 (Light red indicates that the mass-normalized mean values trend higher in the condition above the line, and light yellow indicates that they trend lower). **I.** A schematic illustrating transportation of fatty acids into mitochondria before metabolism. After crossing the cell membrane, free fatty acids are conjugated with coenzyme-A (CoA) to form fatty acyl-CoAs, which are subsequently combined with carnitine. The resulting acyl carnitines can cross the mitochondrial membrane, after which carnitine is released, and the newly-formed fatty acyl CoAs proceed to β-oxidation (created with BioRender.com).

### Metabolic changes caused by stroke

There was also a prominent alteration in the abundance of long chain fatty acids in the ipsilateral hemispheres of stroked mice compared to age-matched naïve controls. By 2 weeks after stroke, the levels of multiple long chain fatty acids were significantly increased, peaking at 4 weeks after stroke; however, the levels were indistinguishable from naïve controls by 12 weeks after stroke (**Figure 3A**). Specifically, there was an increase in palmitate (16:0), eicosenoate (20:1), eicosapentaenoate (EPA; 20:5n3), and docosahexaenoate (DHA; 22:6n3) after stroke (**Figures 3B-E)**. In addition, levels of acyl carnitines, including pentadecanoylcarnitine (C15), oleoylcarnitine (C18:1), docosapentaenoylcarnitine (C22:5n3), and 3-hydroxyhexanoylcarnitine (1), were increased by stroke (**Figure 3F-J**); however, acyl carnitine levels were largely normalized by 8 weeks after stroke. These results indicate that there is an increase in the β-oxidation of fatty acids for at least 4 weeks in the peri-infarct region, which may be due to the catabolism of myelin lipid debris that occurs in the weeks after stroke.^11,12^

**Figure 3.**
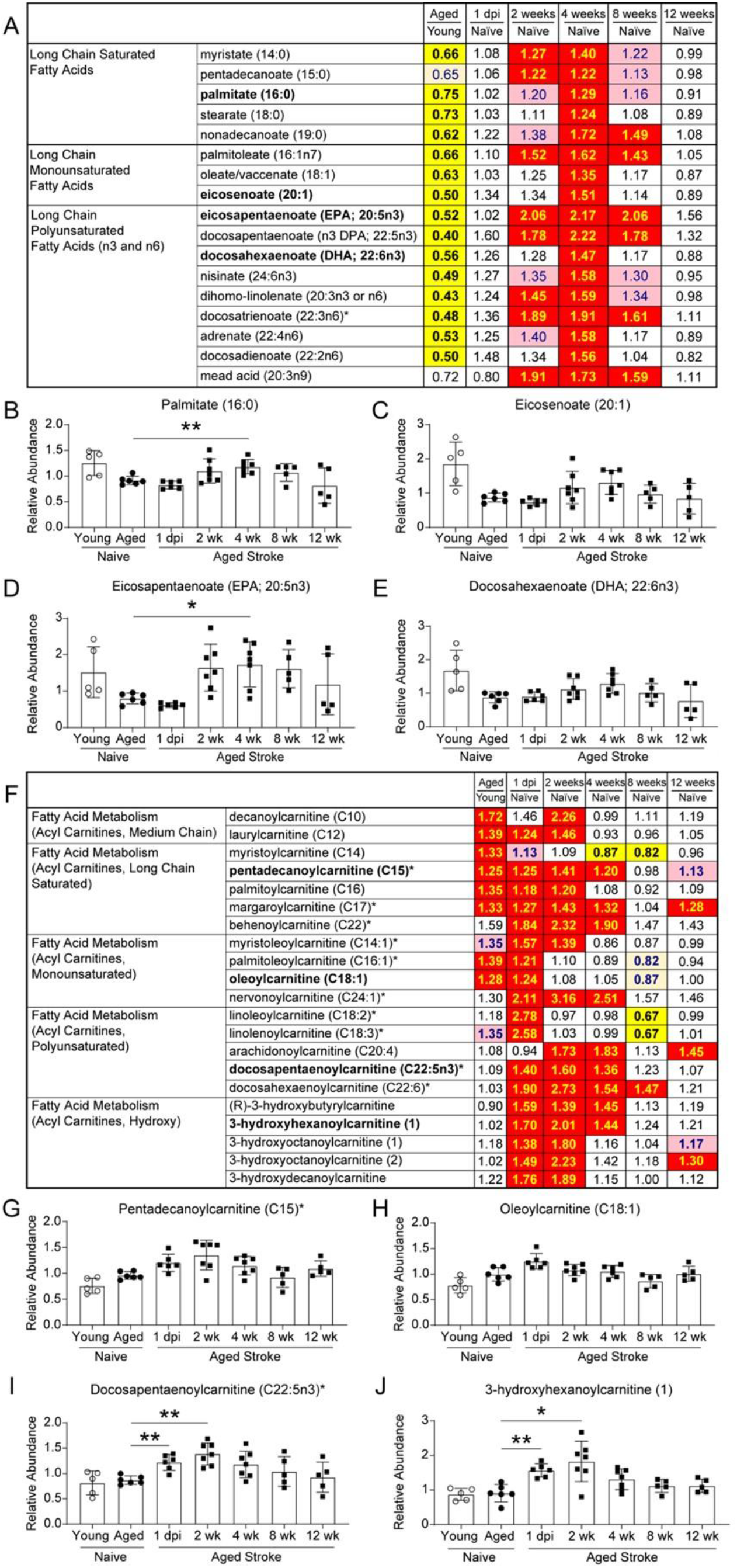
There is an increase in long chain fatty acids and acyl carnitines in the peri-infarct area for at least 8 weeks post stroke. Long chain fatty acids (heatmap, **A**; representative compounds, **B-E**) peaked at 4 weeks after stroke, and several acyl carnitines (heatmap, **F**; representative compounds, **G-J**) were already elevated at 1 day post ischemia. **B**. Palmitate (16:0); Aged naïve vs. 4 wk *\*\*p* = 0.0092. **C**. Eicosenoate (20:1). **D**. Eicosapentaenoate (EPA; 20:5n3); Aged naïve vs. 4 wk \**p* = 0.0308. **E**. Docosahexaenoate (DHA; 22:6n3). **G**. Pentadecanoylcarnitine (C15). **H**. Oleoy-lcarnitine (C18:1). **I**. Docosapentaenoylcarnitine (C22:5n3); Aged naïve vs. 1 dpi \*\**p* = 0.0069, Aged naïve vs. 2wk \*\**p* = 0.0022. **J**. 3-hydroxyhexanoylcarnitine (1); Aged naïve vs. 1 dpi \*\**p* = 0.0036, Aged naïve vs. 2wk \**p* = 0.0287. *n* = 5-7. See Figure 2 for the key to the heatmaps. For the graphs, the groups were compared using Brown-Forsythe and Welch’s ANOVA test, followed by Dunnett’s T3 multiple comparisons test.

Neurotoxic astrocytes secrete toxic phosphatidylcholines (PCs).^13^ Although there were few substantial changes in the expression levels of PCs (**Figure 4A-B**), we found that LysoPCs were markedly elevated for 12 weeks after stroke (**Figure 4A, C**). During inflammation, PCs are converted into LysoPCs by phospholipase A2 (**Figure 4D**), suggesting that the increase in LysoPCs may be caused by chronic inflammation in the peri-infarct region. Phospholipase A2 also leads to the production of pro-inflammatory eicosanoids.^14,15^ Several eicosanoids, including prostaglandin E2, prostaglandin F2alpha, and 12-hydroxyheptadecatrienoic acid (12-HHTrE), were elevated at 4 and 12 weeks after stroke (**Figure 4E**).

**Figure 4.**
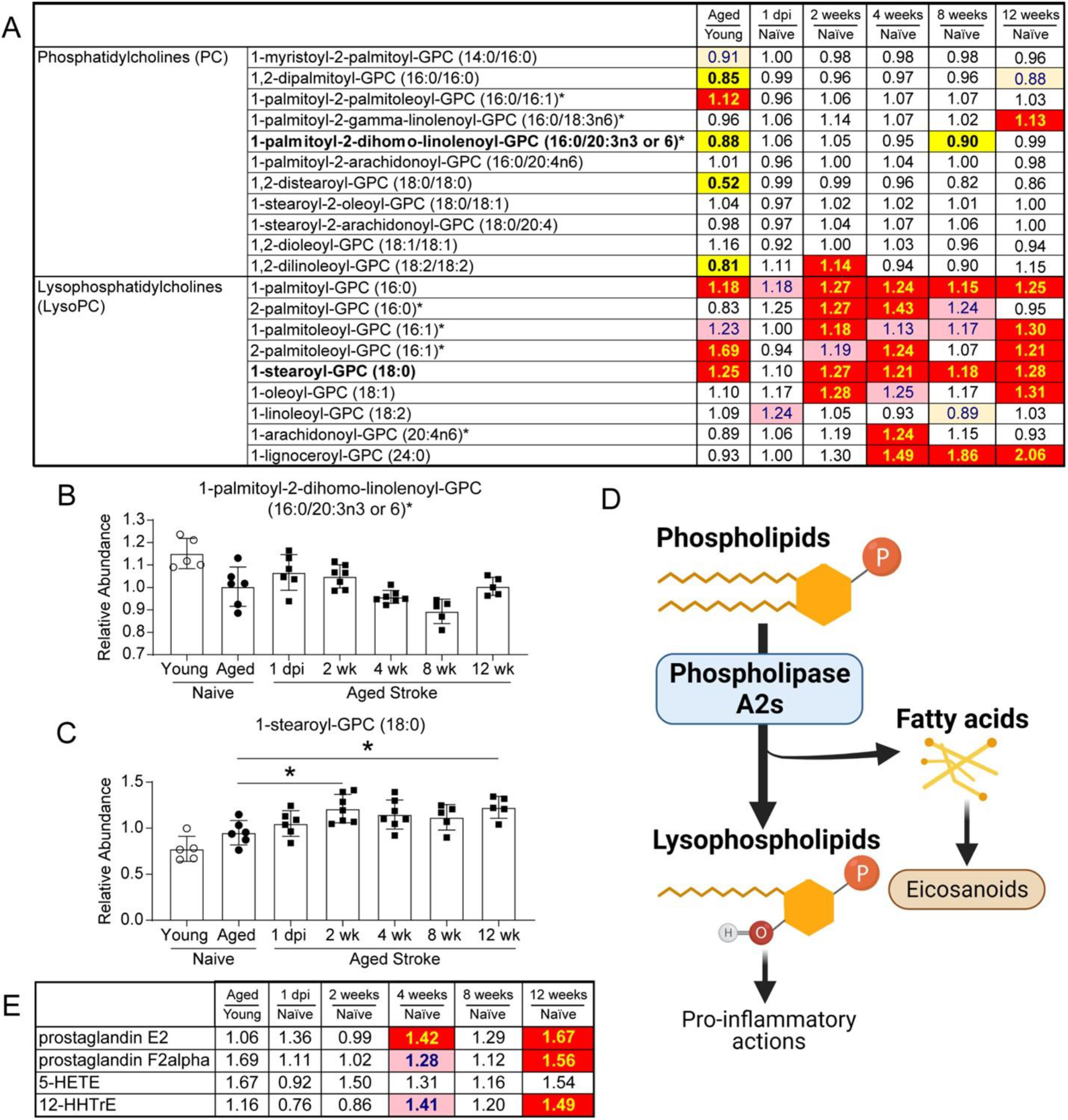
Lysophosphatidylcholines, but not phosphatidylcholines, were increased from 2 weeks after stroke. Lysophosphatidylcholines (**A**; representative compound, **B**), but not phosphatidylcholines (**A**; representative compound, **C**), were increased from 2 weeks after stroke. **B**. 1-palmitoyl-2-dihomo-linolenoyl-GPC (16:0/20:3n3 or 6). **C**. 1-stearoyl-GPC (18:0); Aged naïve vs. 2wk \**p* = 0.0392, Aged naïve vs. 12wk \**p* = 0.0291. *n* = 5-7. **D.** Schematic illustration of the generation of lysophospholipids from phospholipids by Phospholipase A2s (created with BioRender.com). Free fatty acids from this reaction can be turned into eicosanoids (**E**), three of which were elevated at 4 and 12 weeks after stroke. See Figure 2 for the key to the heatmaps. For the graphs, the groups were compared using Brown-Forsythe and Welch’s ANOVA test, followed by Dunnett’s T3 multiple comparisons test.

Glucose levels were also increased in the ipsilateral hemispheres of aged mice after stroke, along with several other glycolysis pathway metabolites (**Figure 5**). However, rather than an increase in glycolysis, we interpret these alterations as the result of a blockage in glycolytic flux because pyruvate, the product of glycolysis, was decreased in the stroke tissue compared to the equivalent tissue from aged naïve mice (**Figure 5B-E**). At 1dpi, an increase in phosphoenolpyruvate (PEP) corresponded with an increase in itaconate. Two weeks post stroke and thereafter, the reduction in pyruvate corresponded with increased dihydroxyacetone phosphate (DHAP) (**Figure 5B, D**). Notably, itaconate is an inhibitor of glycolysis as well as protective against inflammation and oxidative stress (**Figure 5G**).

**Figure 5.**
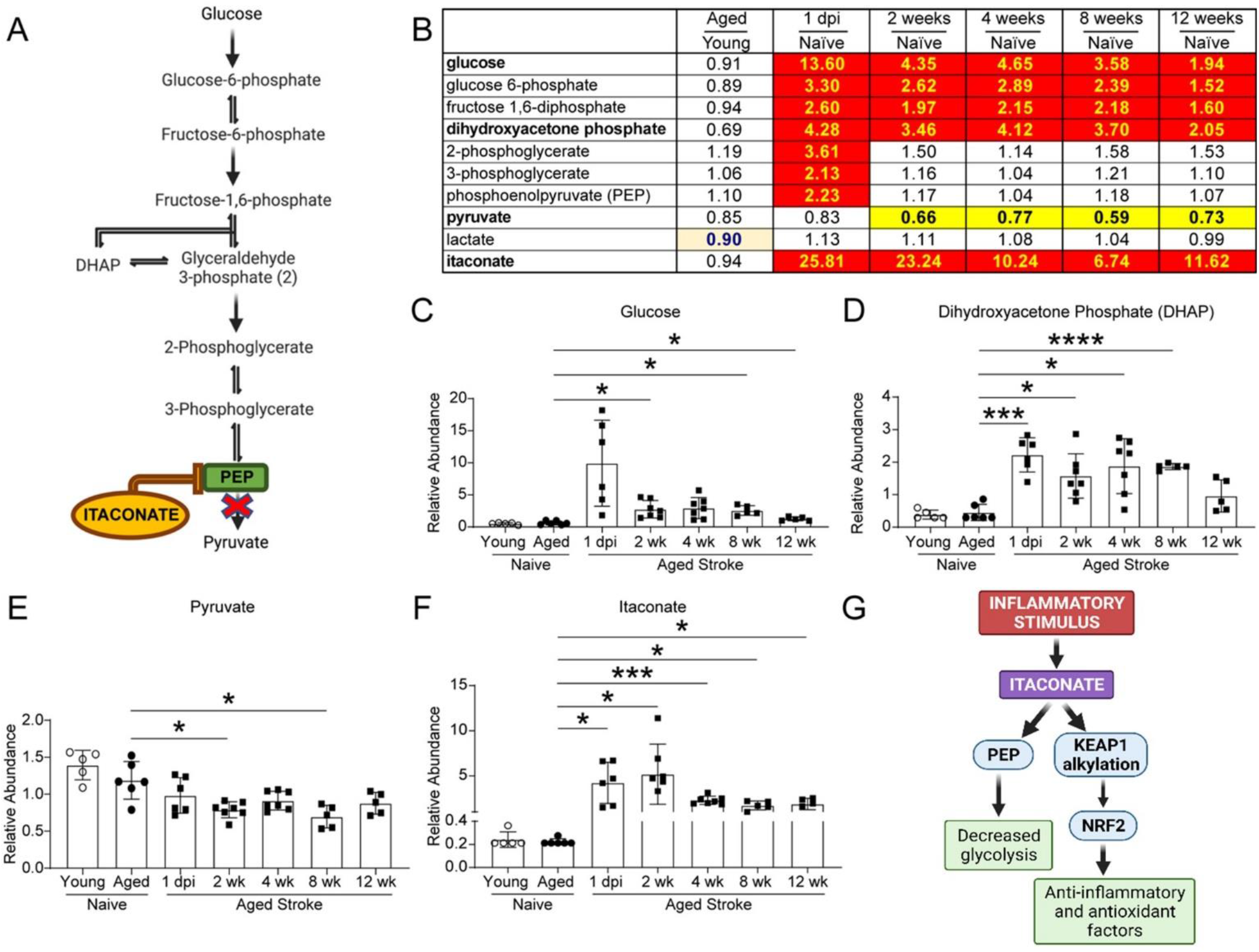
Stroke induces long-lasting alterations in glycolysis and itaconate production. **A-B.** Stroke induces alterations in glycolysis, with some of the changes persisting for at least 12 weeks. **C.** Graphs depicting the levels of glucose; Aged naïve vs. 1 dpi *p* = 0.0850; Aged naïve vs. 2wk \**p* = 0.0252; Aged naïve vs. 4wk *p* = 0.0501; Aged naïve vs. 8wk \**p* = 0.0136; Aged naïve vs. 12wk \**p* = 0.0381. **D.** Dihydroxyacetone phosphate (DHAP); Aged naïve vs. 1 dpi \*\*\**p* = 0.0008; Aged naïve vs. 2wk \**p* = 0.0193; Aged naïve vs. 4wk \**p* = 0.0194; Aged naïve vs. 8wk \*\*\*\**p* < 0.0001. **E.** Pyruvate; Aged naïve vs. 2wk \**p* = 0.0463; Aged naïve vs. 8wk \**p* = 0.0226. **F.** Itaconate; Aged naïve vs. 1 dpi \**p* = 0.0352; Aged naïve vs. 2wk \**p* = 0.0374; Aged naïve vs. 4wk \*\*\**p* = 0.0001; Aged naïve vs. 8wk \**p* = 0.0107; Aged naïve vs. 12wk \**p* = 0.0174. *n* = 5-7. **G**. Itaconate signaling pathway. In response to an inflammatory signal, microglia and macrophages produce itaconate, which is a competitive inhibitor of phosphoenolpyruvate (PEP). Itaconate is also proposed to protect against inflammation and oxidative stress by alkylating Kelch-like ECH-associated protein 1 (KEAP1), thus freeing nuclear factor erythroid 2–related factor 2 (Nrf2) to mediate its multiple anti-inflammatory and antioxidant actions (image created with BioRender.com). See Figure 2 for the key to the heatmap. For the graphs, the groups were compared using Brown-Forsythe and Welch’s ANOVA test, followed by Dunnett’s T3 multiple comparisons test

Regarding neurotransmitters, dopamine, serotonin, and acetylcholine were increased in the aged naïve brain, and glycine and serine were reduced (**Figure 6A, C-D**). No marked changes were seen in glutamate levels (**Figure 6A, B**). However, stroke decreased the levels of glutamate (**Figure 6A-B**), dopamine (**Figure 6A, C**) and adenosine (**Figure 6A, E**) in the ipsilateral hemispheres of aged mice, and the reduction was still present at 12 weeks. The reduction in neurotransmitters after stroke correlated with an increase in neurofilament light in the plasma at 1 day and 2 weeks after stroke (**Figure 6F**).

**Figure 6.**
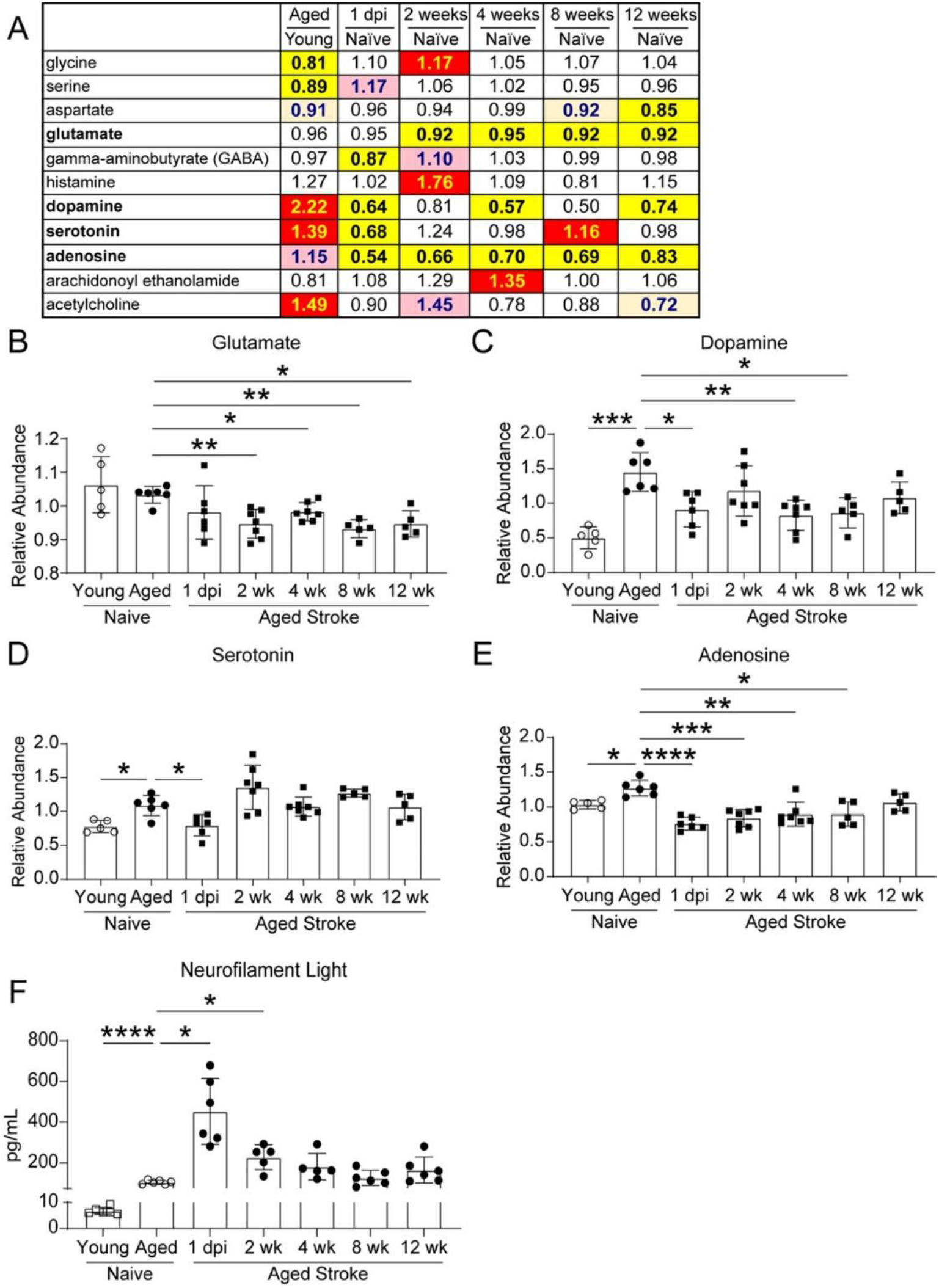
Levels of glutamate, dopamine, and adenosine were reduced for 12 weeks after stroke, and neurofilament light levels in the plasma were increased for 2 weeks after stroke. Heatmap **(A)** and graphical illustrations **(B-E)**. **B**. Glutamate; Aged naïve vs. 2wk *\*\*p* = 0.0064; Aged naïve vs. 4wk \**p* = 0.0283; Aged naïve vs. 8wk \*\**p* = 0.0012; Aged naïve vs. 12wk \**p* = 0.0191. **C**. Dopamine; Aged naïve vs. young naïve \*\*\**p* = 0.0006; Aged naïve vs. 1 dpi \**p* = 0.0315; Aged naïve vs. 4wk \*\**p* = 0.0093; Aged naïve vs. 8wk \**p* = 0.0196. **D**. Serotonin; Aged naïve vs. young naïve \**p* = 0.0153; Aged naïve vs. 1 dpi \**p* = 0.0413. **E**. Adenosine; Aged naïve vs. young naïve \**p* = 0.0103; Aged naïve vs. 1 dpi \*\*\*\**p* < 0.0001; Aged naïve vs. 2wk \*\*\**p* = 0.0002; Aged naïve vs. 4wk \*\**p* = 0.0042; Aged naïve vs. 8wk \**p* = 0.0214, *n* = 5-7. **F**. Neurofilament Light; Aged naïve vs. young naïve \*\*\*\**p* < 0.0001; Aged naïve vs. 1 dpi \**p* = 0.0156; Aged naïve vs. 2wk \**p* = 0.0475. *n* = 5-7. See Figure 2 for the key to the heatmap. For the graphs, the groups were compared using Brown-Forsythe and Welch’s ANOVA test, followed by Dunnett’s T3 multiple comparisons test.

## Discussion

Little is currently known about how metabolism changes with aging in the rodent brain despite many factors likely contributing to age-dependent changes in brain metabolism. For example, changes in the blood-brain barrier (BBB)^16^ may result in the accumulation of plasma proteins, which can cause inflammation and interfere with neuronal circuitry.^17^ Additionally, alterations in mitochondrial structure and function have been recognized as a hallmark of the aging brain.^18,19^

Recently, however, a group led by O. Fiehn composed a metabolome atlas of the aging mouse brain at 3 weeks, 4 months, 15 months, and 2 years.^20^ They found that several acyl carnitines, like myristoylcarnitine (C14), palmitoylcarnitine (C16) and palmitoleoylcarnitine (C16:1) were increased with age, while free saturated fatty acids were decreased. They also found an increasing trend in levels of dopamine and acetylcholine in 15 mo mice, when compared to the 4 mo mice. The results obtained from our comparative study between young adult and middle-aged mice are in line with these findings.

However, there are also differences between our study and this investigation, specifically related to the levels of several long chain fatty acids. For example, Ding et al. found that the levels of eicosenoate and docosahexaenoate (DHA) stayed the same instead of decreasing between 4 months and 15 months of age. These differences may arise from the fact that the mice used in our study were from 7 months to 18-20 months of age rather than 4 to 15 months. Moreover, the brain metabolomic atlas generated by Ding et al. shows that levels of metabolites can differ drastically between brain regions, and our samples represent mean values from the ipsilateral hemisphere of the brain, which consists of cortex, hippocampus, thalamus, and basal ganglia regions in our stroke model. We were unable to divide the brain into smaller regions for this study due to the amount of tissue required for our unbiased global metabolomic analysis.

Neurotransmitters, such as dopamine and serotonin, decline with aging in the human brain, but in rodents, the reduction has been shown to be smaller and more variable.^21,22^ We observed an increase in these neurotransmitters when we compared young naïve mice to aged naïve mice, despite an increase in neurofilament light in the plasma of aged mice. The reason for this discrepancy may be that the age of our older cohort was sufficient to cause age-related neurodegeneration and increase neurofilament light in plasma, but not sufficient for compensatory mechanisms to prevent a decline in neurotransmitter levels.

Little is also currently known about the metabolic changes that occur in the brain in the recovery period after stroke. To our knowledge, existing studies have been restricted to the brains of young rodents and only at acute (1-3 days), and sub-acute (7 days) time points post stroke.^23–27^ Age is the largest non-modifiable risk factor for ischemic stroke,^1^ and metabolic changes in the aging brain may make it more vulnerable to the aftereffects of stroke. Therefore, in this study, we also evaluated the metabolic changes that occur in the brain in aged mice at acute, subacute, and chronic time points.

We found that long chain fatty acids were elevated between 2 and 8 weeks after stroke, and LysoPCs were elevated for 12 weeks after stroke. These alterations may be the result of myelin degradation, as we have previously shown that there is a similarly delayed accumulation of cholesterol crystals, sphingomyelins, sulfatides, and lipid droplets after stroke.^11,12^ However, a group led by S. Liddelow^13^ recently reported that neurotoxic reactive astrocytes, induced by the secretion of Il-1α, tumor necrosis factor alpha (TNF-α), and C1q by activated microglia,^28^ induce cell death via the release of long chain saturated fatty acids and PCs. Fatty acids act as important secondary messengers in cell signaling, but they can also have pro-inflammatory effects,^29^ and the accumulation of fatty acids in the brain has been associated with several neurodegenerative disorders.^30^ Therefore, neurotoxic reactive astrocytes in the peri-infarct region could also be the source of these lipids. Future studies are necessary to determine the source of the increase in long chain fatty acids.

Alongside the increase in fatty acid levels, we found that there was an increase in acyl carnitine levels. These alterations indicate that the long chain fatty acids may be undergoing β-oxidation for the formation of acetyl-CoA as fuel for the TCA cycle because long chain fatty acids destined for oxidation are conjugated to carnitine by the outer mitochondrial membrane carnitine-palmitoyltransferase 1 (Cpt1), which results in the release of coenzyme A (CoA). The resulting acyl carnitines are then transported across the intermembrane space by carnitine/acylcarnitine translocase and acted upon by the inner membrane carnitine-palmitoyltransferase 2 (Cpt2), which releases carnitine and reconjugates the long chain fatty acid with CoA for mitochondrial fatty acid β-oxidation. This could be advantageous for the brain for two reasons: 1) it provides an alternative source of acetyl-CoA in light of our finding that glycolytic flux appears to be reduced, and 2) it provides a method of catabolizing fatty acids derived from myelin breakdown or produced from neurotoxic reactive astrocytes. However, high levels of acyl carnitines and LysoPCs have also been associated with mitochondrial dysfunction and inflammation after ischemia.^31,32^ Future studies are also required to determine the role of increased fatty acid oxidation after stroke.

Fat occupies approximately 60% of the dry weight of the brain, making it the most lipid-rich organ in the body.^33^ Despite the large fat content, the brain predominantly uses glucose for energy, with the utilization of fatty acids estimated to be up to 20% of the total energy consumption of the brain.^34^ Our results demonstrated that many glycolysis-related metabolites were upregulated at 1 day after stroke and were still elevated at 12 weeks. Concurrently, pyruvate was decreased, and itaconate, a glycolysis inhibitor, was found to be increased. Itaconate has been proposed to disrupt glycolysis at the level of pyruvate kinase via competitive inhibition due to its structural similarity to PEP.^35^ These data indicate that glycolytic flux is impaired for at least 12 weeks after stroke and raise the possibility that itaconate is an early regulator of this alteration in metabolic homeostasis. Notably, the increase in PEP may be another level of regulation, as PEP is a competitive inhibitor of the enzyme triosephosphate isomerase (TPI), which converts DHAP into glyceraldehyde 3-phosphate. In support of this possibility, DHAP levels were increased for 12 weeks after stroke.^36^ Therefore, glycolysis may initially be inhibited after stroke at the level of pyruvate kinase via the accumulation of itaconate and also at the level of TPI via the accumulation of PEP.

Itaconate is an immunometabolite discovered in 1840 by G. Crasso.^37^ However, its role in mammalian metabolism was unclear until 2011, when it was first reported to participate in mammalian immune responses.^38–40^ Itaconate is produced in microglia and macrophages by the action of the highly inducible gene *aconitate decarboxylase 1* (*Acod1*, also known as *Irg1*; *Immunoresponsive gene 1*) when the phagocytes respond to an inflammatory stimulus, such as an infection or injury. The enzyme produced by *Acod1* catalyzes the production of itaconate from the TCA cycle metabolites citrate and cis-aconitate. Importantly, the induction of itaconate can protect against oxidative stress and is anti-inflammatory via the alkylation of kelch-like ECH-associated protein 1 (KEAP1).^41^ Under normal conditions, KEAP1 exists to inhibit nuclear factor erythroid 2–related factor 2 (Nrf2) in the cytoplasm; however, alkylation of KEAP1 leads to accumulation of Nrf2, resulting in multiple antioxidant and anti-inflammatory actions (**Figure 5G**).^42,43^

Inflammation, gliosis, and immune cell infiltration into peri-ischemic regions of the ipsilateral hemisphere results in an increase in microglia, astrocytes, and peripheral immune cells compared to control tissues. This difference in cellular composition of the ipsilateral hemisphere almost certainly contributes to the metabolic differences reported in this study. Nevertheless, to the best of our knowledge, this study is the first global characterization of the metabolic profile of the aging mouse brain during the first 12 weeks of recovery following ischemic stroke and still provides novel insight into the changes in metabolic homeostasis that occur in the brain during recovery. The results of this study provide a foundation for future studies focused on determining the underlying functions and cell specificity of these changes in metabolic homeostasis.

In conclusion, this study has revealed that stroke alters glycolysis and lipid metabolism in the brain for at least 12 weeks. The brain is highly reliant on glucose for energetic requirements. Although fatty acid oxidation does occur within the brain, it is slower and consumes more oxygen than ATP generation from glucose and exposes neural cells to more oxidative stress.^44^ The changes in metabolism could reflect 1) mitochondrial dysfunction due to brain tissue damage caused by injury and the resulting inflammatory response, 2) adaptation to altered fuel abundance, or 3) differing cell populations in the recovering hemisphere compared to control tissues. This study also revealed that stroke chronically reduces levels of neurotransmitters in the surviving ipsilateral tissue and raises the possibility that the immunometabolite itaconate may be a central regulator of the chronic inflammatory response to stroke via the metabolic reprogramming of myeloid cells.

## Acknowledgements

This work was funded by NINDS R01NS096091 (KPD), NIA R01AG063808 (KPD), United States Department of Veterans Affairs I01RX003224 (RGS), and the Leducq Foundation Transatlantic Network of Excellence Stroke-IMPaCT (KPD). We are also grateful to Metabolon Inc. for their expertise and assistance throughout all aspects of our study.

## Author contribution statement

SL, MTG, BKM, KEJ, and KD conducted the research, SL, DB, KD, and RS wrote the manuscript.

## Supplementary material

**Supplementary Figure 1.** Principal component analysis of controls.

**Supplementary Table 1:** The number of significantly different metabolites for each comparison

**Supplementary Table 2:** The fold change and *p* value for the detected metabolites for each comparison

