## Supplementary Figure 1 for "Increased fatty acid metabolism and decreased glycolysis are hallmarks of metabolic reprogramming in the brain during recovery from experimental stroke"

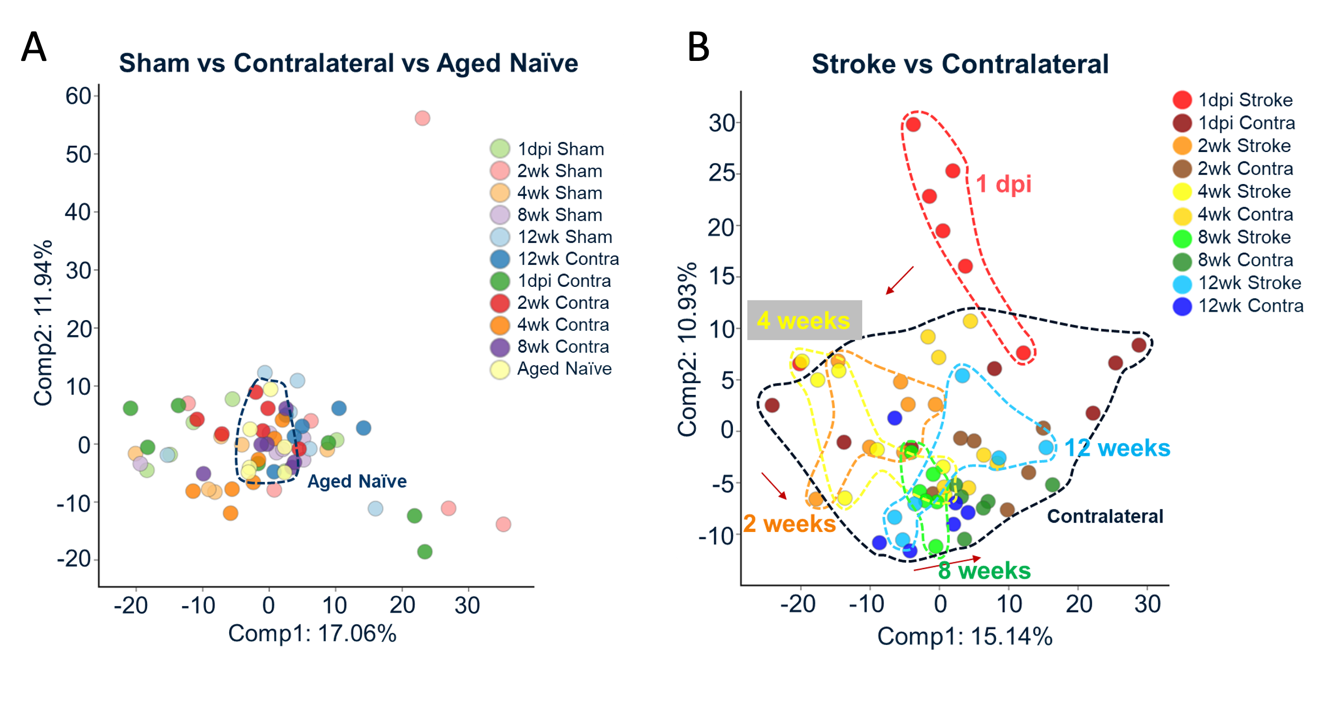

**Supplementary Figure 1. Principal component analysis of controls.** **A.** Despite some differences between the different control groups, sham mice and the contralateral hemispheres from the stroked mice still cluster close to the aged naïve controls. **B.** There is still separation based on stroke time point when using the contralateral hemispheres of the stroked mice as a control instead of the aged naïve mice used in **Figure 1D**.
