## Supplementary Table 1 for "Increased fatty acid metabolism and decreased glycolysis are hallmarks of metabolic reprogramming in the brain during recovery from experimental stroke"

**Supplementary Table 1. Number of significantly differed metabolites in each of the comparisons.** Blue indicates the number of metabolites increased, and red indicates the number of metabolites decreased in the condition above the line.

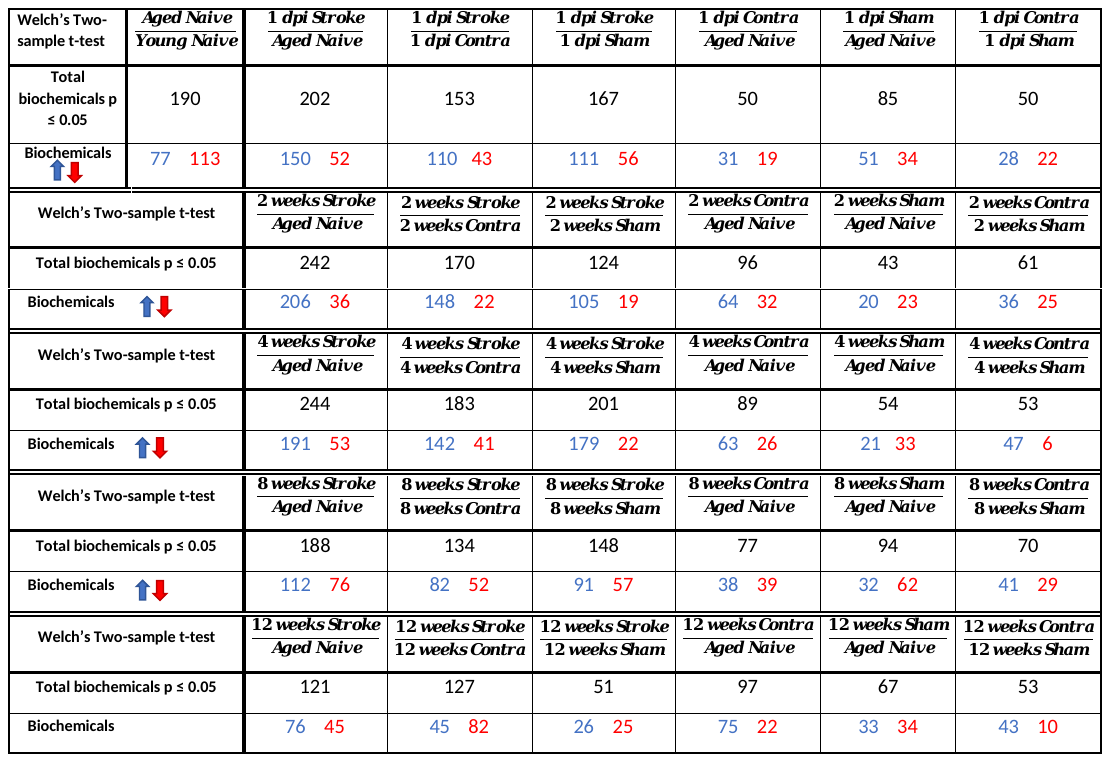

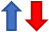
